# Intraventricular hemodynamics in pediatric patients with single right ventricles reveal deteriorated washout and low vortex formation times: An in silico study

**DOI:** 10.1101/2020.12.13.422573

**Authors:** Anna Grünwald, Jana Korte, Nadja Wilmanns, Christian Winkler, Katharina Linden, Ulrike Herberg, Sascha Groß-Hardt, Ulrich Steinseifer, Michael Neidlin

## Abstract

The congenital heart disease univentricular heart (UVH) occurs with an incidence of 0.04-0.5% in newborns and is often treated with the Fontan procedure. In this intervention, the cardiac circulation is transformed into a singular circulation with only one ventricular chamber pumping.

Hemodynamics the singular ventricle is a major research topic in cardiology and there exists a relationship between fluid dynamical features and cardiac behavior in health and disease. By visualizing the flow using Computational Fluid Dynamics (CFD) models, an option is created to investigate the flow in patient-specific geometries.

CFD simulation of the pathological single right ventricle in contrast to the healthy left ventricle is the research object of the present work. The aim is the numerical comparison of the intraventricular flow within the ventricles. Based on this, flow formation in different anatomies of the ventricles is investigated.

Patient-specific measurements of ventricles from three-dimensional real-time echocardiographic images served as the basis for the simulations with five single right ventricle (SRV) patients and two subjects with healthy left hearts (LV) investigated. Interpolation of these data reproduced the shape and continuous motion of the heart during a cardiac cycle. This motion was implemented into a CFD model with a moving mesh methodology. For comparison of the ventricles, the vortex formation as well as the occurring turbulent kinetic energy (TKE) and washout were evaluated. Vortex formation was assessed using the dimensionless vortex formation time (VFT).

The results show significantly lower values for the VFT and the TKE in SRV patients than for the compared LV Patients. Furthermore, vortex formation does not progress to the apex in SRV patients. These findings were confirmed by a significantly lower washout in SRV patients.

Flow simulation within the moving ventricle provides the possibility of more detailed analysis of the ventricular function. Simulation results show altered vortex formation and reduced washout of SRV in comparison to healthy LV. This information could provide important information for the planning and treatment of Fontan patients.

## Introduction

The heart defect single ventricle syndrome describes those patients who are born with only one “functional” heart chamber (univentricular heart (UVH)) [1]. There is a probability of 0.04-0.5% that newborns have a univentricular heart. When pediatric patients are found to have this type of cardiac defect, it must be treated within the first few days after birth by regulating the blood supply to the lungs [2].

In the course of the first two years of the patient’s life, surgical intervention is performed with the Fontan procedure, which enables the body’s circulatory system to be supplied with only one ventricle. Oxygen-depleted blood is directed to the lungs instead of the heart, and the one functional ventricle takes over the circulation of oxygen-rich blood through the body [3].

The Fontan procedure is a successful palliative strategy for patients with single-ventricle syndrome. However, its inherent limitations and deficits are becoming apparent as more patients present with failing Fontan physiology [4]. Flow properties in the ventricle provide information about the health of the cardiovascular system. Consequently, their study can be used to assess the function of the ventricle [5].

Flow within the ventricle is strongly influenced by cardiac motion and shape [6]. In this study, we will compare the intraventricular flow within the left ventricles (LV) from patients with healthy hearts and pathological single right ventricles (SRV) from Fontan patients. The comparability is based on the work to be done by the SRV in the systemic circulation, which is equivalent to the work done by the LV [7].

A numerical flow analysis is conducted to enable the comparison of the intraventricular flows. For this analysis a moving mesh model is used, which incorporates the geometry changes of the ventricular walls as a boundary condition in the simulation. The simulation uses segmented and time-discrete echocardiographic images from which the patient specific 3D geometry is extracted. Through interpolation between the acquired data images, a continuous cardiac motion is represented.

## Methods

### Data acquisition

Volume-time curves of five patients with single right ventricle (SRV) circulation and two subjects with a healthy left ventricle (LV) were acquired retrospectively from 3D echocardiography measurements performed at the University Hospital Bonn (UKB). The ultrasound device iE33 (Philips Medical Systems, Andover, US) was used to record the volumetric shape of the ventricle over one cardiac cycle. Between 12-28 steps per heartbeat were recorded. The study was approved by the local ethics institutional review committee (Registration No. 226/06) and conformed to the principles of the Declaration of Helsinki as well as German law.

The SRV group was selected in order to represent different ages, end diastolic volumes (EDV) and ejection fractions (EF). The LV group was selected in order to have similar ages, EDVs and EF>55%. Table 1 summarizes the patient data used in this study.

**Table 1:** Clinical data for univentricular heart (SRV) and healthy heart (LV) patients (EDV: End diastolic Volume, bpm: beats per minute, EF: ejection fraction)

| # | Ventricle | Gender | Age (yrs) | EDV (ml) | bpm | EF (%) |
| --- | --- | --- | --- | --- | --- | --- |
| 1 | SRV | m | 6 | 26 | 93 | 45 |
| 2 | SRV | m | 8 | 28 | 79 | 52 |
| 3 | SRV | m | 6 | 55 | 93 | 51 |
| 4 | SRV | f | 17 | 97 | 60 | 45 |
| 5 | SRV | m | 15 | 100 | 59 | 39 |
| 6 | LV | m | 4 | 40 | 153 | 67 |
| 7 | LV | f | 7 | 54 | 98 | 58 |

The geometric data from the echocardiography was translated into a unicode character database (UDC) format using the program ImageArena (TOMTEC Imaging Systems, Unterschleißheim, Germany). For each time step a 3D model with a constant number of cells and associated cell nodes was created. Afterwards, the UCD data was translated using Matlab into an STL file format for the application of the moving mesh model.

For the interpolation between the individual time steps and the integration of the continuous ventricle surface movement, it was necessary that the outer mesh of the recorded geometry ensured a constant number of nodes in each time step and a high-quality mesh. In order to prevent partially occurring problems of the mesh display during the simulation, modifications of the outer surface of the geometry were performed with the open source program Blender 2.82 (Blender Foundation).

### Simulation

The meshing of the geometry was performed with the ANSYS Meshing tool from ANSYS 2019 R1 (Ansys Inc., Canonsburg, US) with unstructured tetrahedral elements supported by a “Patch Conforming Method”. The boundary layer was represented in two cases with 8 and 15 prism layers. Mesh independence was determined based on the area average velocities at the inlets and outlets and the maximal wall shear stress during peak diastolic flow (E-wave) and changes in the variables of <5% between two mesh refinement steps were considered independent. An y+ value of < 5 was taken as a measure for appropriate resolution of the boundary layer flow. Comparing the peak average velocities at the inlet and outlet of the ventricle for each cycle a threshold of <5% was taken. Mesh independent results converged from 750.000 element number.

The mesh model utilized smoothing and layering to ensure appropriate mesh quality during the deformation of the domain. The function “diffusion cell volume based” was used as the smoothing approach to adapt the inner mesh of the geometry to the movement of the outer surface. The diffusion parameter was set to 0, which resulted in a uniform diffusion of mesh deformation. The layering was ratio-based and dynamic, so that the inflation layers also adapted to the movement of the outer surface.

For the simulations the non-Newtonian blood model by Ballyk et al. [8] with a density of ρ = 1056.4 kg/m3 (hematocrit level of 44%) was applied. In addition, the pressure-based solver was selected and set in coupled mode and solved with ANSYS Fluent. The flow was assumed to be turbulent and the kω-SST model was applied. For the convergence of the velocity, k- and ω-residuals the criteria were set to 0.001 and for the continuity to 0.0003. The maximum number of iterations was 30. The number and size of the time steps depended on the frequency of the calculated boundary conditions, as does the number of interpolations. Resulting time steps ranged between 0,0018 and 0,004s. The simulations started in the end systolic state of the cardiac cycle and run for five cycles.

As inlet and outlet conditions, patient specific pressure curves were set as boundary conditions. Patientspecific pressure curves for the srv patients were generated using clinical data from UKB. The pressure curves for the healthy LV were based on calculated standard values. Valves were not modeled and a wall or a fully opened inlet/outlet were set. at the open valve surface and the currently closed valve as a wall.

Ventricular movement was achieved with the moving mesh method. First, a UDF (“AssignID”) for a mesh geometry was written in ANSYS Fluent. For each cell node the corresponding point identification number and the Cartesian coordinates were written to a file (“surface”). In the surface file only the data of the cell nodes of the ventricular wall and the entrance and exit surfaces were stored.

In the second step, the mesh of the original STL files (see above) was linked to the cell nodes created and stored in ANSYS (surface-file). This was done via a script created with Python. First, for each cell node that is read out, the corresponding area on the STL file and the position of the node on this triangular area was determined. For this purpose, parameters (α and β) were set up which describe the position of the point on the area in relation to the three nodes (P1, P2 and P3) of the triangular area (equation (2) For each cell node the calculated values of α and β were stored in a file together with the identification number of the point.

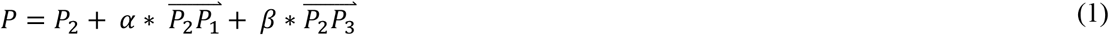

The stored values were used to determine the position of the cell nodes on the subsequent STL files and a “surface_n” file was created for each existing STL file. This file contains the point identification numbers with the corresponding x,y,z coordinates of the node.

In the third step, the temporal interpolation between the mesh data was performed to enable continuous movement in the simulation. With another script written in Python, interpolation functions for the movement were calculated for each cell node. Subsequently the regularly interpolated points between the respective cell nodes were calculated and read out into the “udfsurface_n” files. As interpolation method the cubic spline interpolation was used. The interpolation steps corresponded to the time step size in the CFD simulation (see above). In the last step, the calculated continuous mesh movement was loaded into the simulation.

### Flow analysis

For the evaluation of the blood flow characteristics, the vortex structure formation, the vortex formation time, turbulent kinetic energies and the ventricular washout were investigated.

The vortex structure formation was visualized by the Q-criterion within the ventricle. The Q-criterion describes the local balance between rotation and shear in all vector directions. The calculation was performed using the second invariant of the velocity gradients, which are summarized by the symmetric shear rate tensor S and the antisymmetric vorticity tensor Ω. [9]

The dimensionless parameter vortex formation time (VFT) describes the process of vortex formation during the beginning of diastole. It represents the length of the jet, normalized to the mitral valve diameter. The VFT is calculated as the time integral of the velocity from the start of the diastole to the peak of the E-wave. It measures the quality of the vortex formation process and is a characteristic value for ventricular filling.(Gharib etal.,1998; Kheradvar and Pedrizzetti, 2012).

Turbulent kinetic energy (TKE [J/kg]) was calculated in Ansys Fluent based on the transport equation for the k-ω-SST turbulence model. This model is based on the standard k- ω model from Wilcox [10]. It uses the transport equations for the turbulence kinetic energy (k ≙ TKE [J/kg) ((2)) and the specific dissipation rate (ω). In these equations, G_k_ represents the generation of turbulence kinetic energy due to mean velocity gradients. Г_k_ represent the effective diffusivity of k, respectively. Y_k_ represent the dissipation of k. S_k_ is a user-defined source terms.

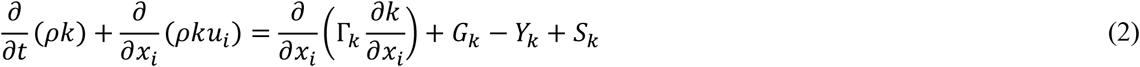

The maximum value of TKE within the ventricle was recorded for each patient over five cycles. To reduce scatter in the data, the last four cycles were superimposed and the median was taken for each time step. From this smoothed cycle, all TKE values were accumulated over the entire cycle. In this way, the total average TKE for each patient could be determined. In addition, the ratio of total systolic TKE to total TKE was examined.

Occurring blood stagnation, which indicates that the ejection performance does not meet the requirements of blood exchange, was evaluated by means of washout simulations as shown in [11]. In brief, simulations were performed with two fluids 1 and 2 with identical rheological properties (see above). In the beginning, the entire domain was filled with fluid 1 (old blood), afterwards the volume fractions were switched and simulations with fluid 2 (new blood) were performed for three cycles. Treatment of both fluids were expressed with the Volume of Fluid methodology. For more details see [11].

The results of the fluid analysis for the SRV group are shown as mean value +- standard deviation. Furthermore, statistical comparison between the SRV group and the individual LV was performed with a one-sample t-test and a significance threshold of p=0.001.

## Results

### Flow description

Figure 1 shows the flow field inside the ventricle for patient (#3) with SRV and patient (#7) with a healthy LV during diastole and systole. Six time points of the cardiac cycle are represented; E-wave, diastase, A-wave, start systole, peak systole, end systole. The blood flow over a cycle and the movement of the vortex is shown through blue isosurfaces based on the Q-value. Furthermore, the TKE is represented in the observed time points. For better visualization purposes different scales for TKE and Q-Value were applied, see (Table 2).

**Figure 1:**
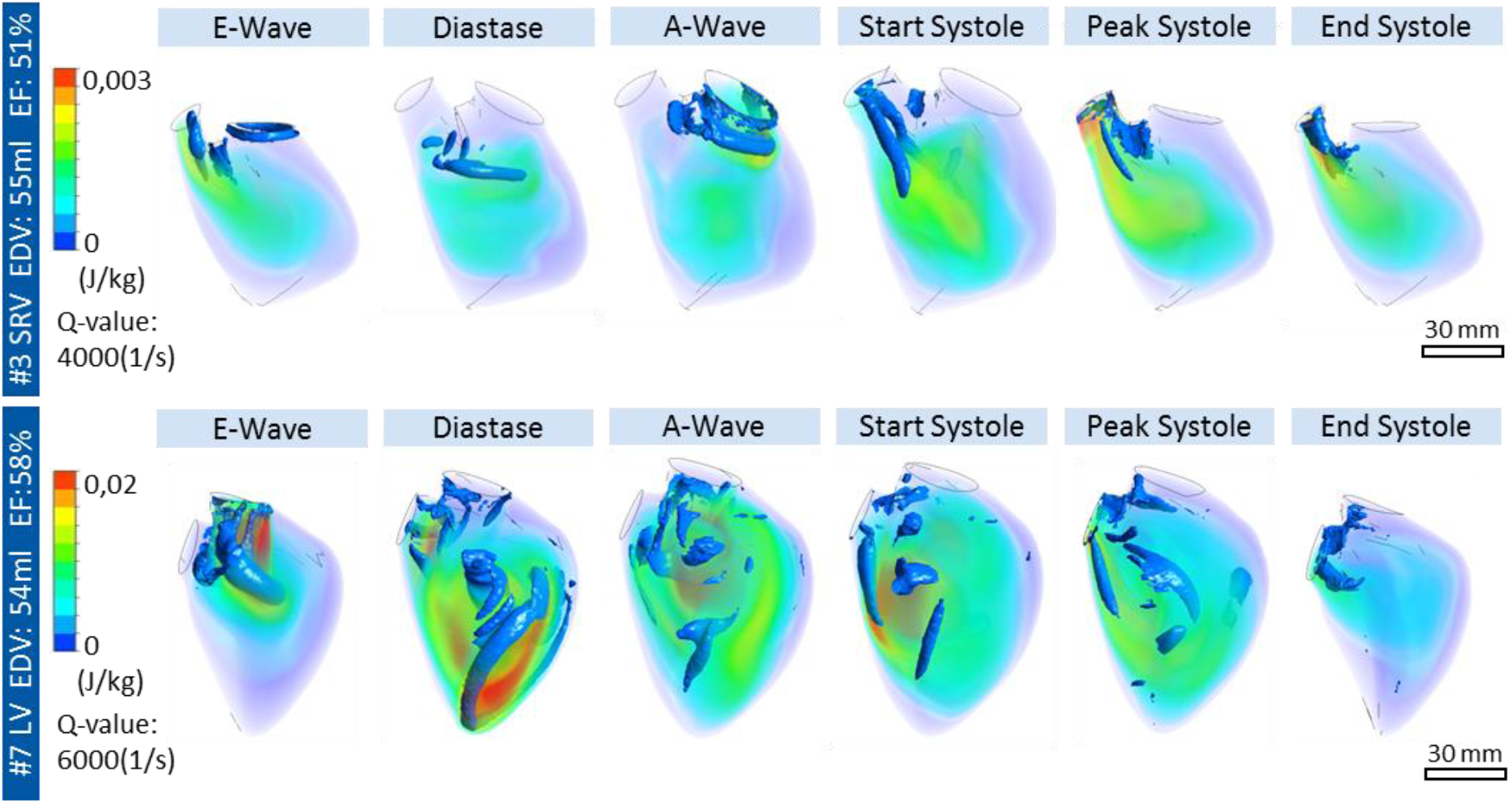
Turbulent kinetic Energy (J/kg) and vortex structure formation with Q-value (1/s) for subject #3 (SRV) and subject #7 (LV)

**Table 2:** Results for single right ventricle (SRV) group (mean value, standard deviation) and left ventricle (LV) patient.

|  | **SRV** | **LV #6** | **P-value** | **LV #7** | **P-value** |
| --- | --- | --- | --- | --- | --- |
| TKE_ges (J/kg) | $0.881 \pm 0.708$ | 23.713 | $<1e-5$ | 5.438 | 0.00014 |
| TKE_sys/ TKE_ges (-) | $0.543 \pm 0.165$ | 0.442 | 0.24313 | 0.386 | 0.09976 |
| VFT (-) | $1.16 \pm 0.53$ | 4.26 | 0.00019 | 3.96 | 0.00028 |
| EF (%) | $46.4 \pm 4.71$ | 58 | 0.00533 | 67 | 0.00062 |
| Washout (%) |  |  |  |  |  |
| 1.Cycle | $43.79 \pm 9.77$ | 63.05 | 0.01162 | 57.43 | 0.03544 |
| 2.Cycle | $64.86 \pm 9.54$ | 87.26 | 0.00630 | 85.54 | 0.00836 |
| 3.Cycle | $76.32 \pm 6.64$ | 95.51 | 0.00296 | 94.88 | 0.00335 |

The presented SRV of subject #3 has an EDV of 55 ml and an EF of 51%. The beginning of the E- and A-wave can be recognized by a developing vortex at the tricuspid valve (inlet). During diastasis, the E-wave vortex only propagates in the upper third of the ventricle. The second developing vortex of the A-wave propagates comparably. During systole, an elongated vortex develops in the central part of the ventricle and is ejected at the end of systole. The presented SRV-case is exemplary for all five examined SRV-patients where a similar vortex progression can be observed. In all cases the vortex formation does not extend to the apex of the ventricle.

The exemplary blood flow in a healthy ventricle is also shown in Figure 1. The healthy subject has an LV with an EDV of 54 ml and an EF of 58%. During the E-wave, a circular vortex is formed starting from the mitral valve. During diastasis, the vortex and the TKE maximum move towards the apex. During A-wave, the vortices are mixed and scattered eddies can be observed. At the beginning of systole, elongated vortices migrate to the outlet and are ejected until the end of systole.

TKE is almost an order of magnitude higher in the healthy LV in comparison to subject #3. The differences in vortex formation are further underlined by the lower threshold of the Q-criterion to achieve any visualization of the flow structures (6000 s-1 – LV vs. 4000 s-1 - SRV).

### Flow analysis

The results for the flow analysis for five cardiac cycles of each patient are summarized in Table 2.

The TKE of 0,881 ± 0,708 (J/kg) in SRV patients is significantly lower than in the compared healthy patients. Additionally, it is observed that the SRV ratio of systolic TKE to total TKE is slightly higher in SRV patients than in healthy patients. However, in two cases of SRV patients, the ratio was 73% and 75%, respectively.

VFT for the SRV patients was with 1.16 ± 0.53 significantly lower than the compared healthy LV patients. The two healthy LV patients show a VFT of 4.26 and 3.96.

The SRV patients exhibit a significantly lower EF than the LV patients. The mean deviation for the SRV is 4.71% and out of the five SRV patients examined, two patients show an EF of more than 50%.

According to Table 2, the average inflow of new blood (phase 2) in SRV after a cycle is 43.79%. After the second cycle, the percentage for SRV is 64.86% and over the course of three cycles, an average of 76.32% of the new blood is present in the SRV. For LV patients #6 and #7, 95.51% and 94.88% of the new blood is present after three cycles which are significantly higher than the SRV group (p=0.008 and p=0.0033).

The washout characteristics of SRV (patient #3) and healthy LV (patient #7) are shown in Figure 2, left pane. Within three cardiac cycles the time point of diastasis is displayed. The distribution of new blood (phase 2) colored red in the SRV during the first cycle is only in the upper part of the ventricle. This structure continues to develop in the second cycle towards the apex. At the end of the third cycle, however, residual blood (phase 1) colored blue is still visible in the apex. A lack of washout in the apex is observed in all SRV patients. In contrast, during diastole of the LV of the first cycle, new blood is distributed in the area below the valve and in the apex. This distribution can also be seen in the diastase of the second cycle. At the end of the third cycle, the ventricle is 95% filled with new blood.

**Figure 2:**
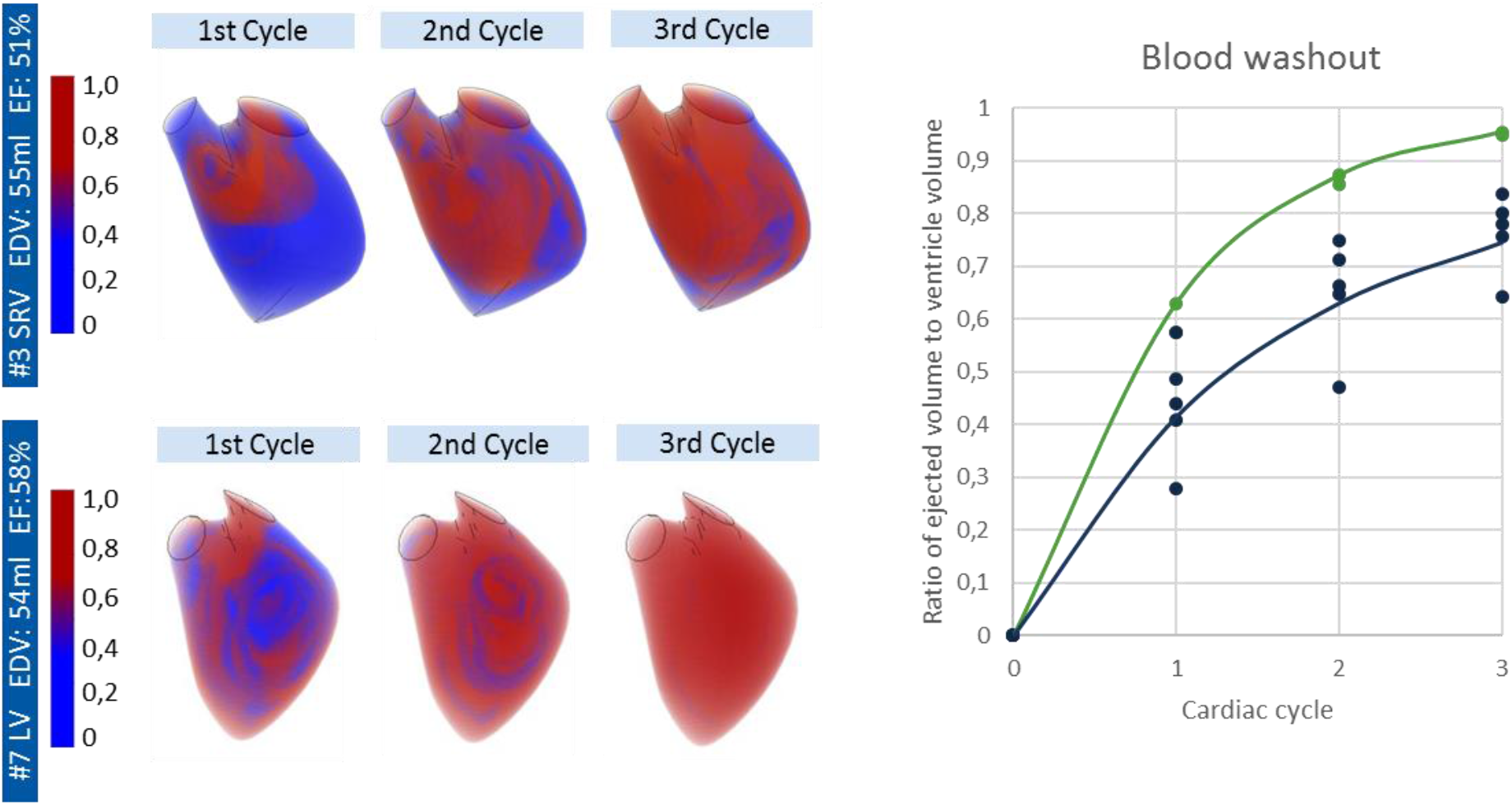
Left: Washout simulation for subject #3 (SRV) and subject #7 (LV). Time point of diastase for tree cardiac cycles. Blue old blood, red new blood. Right: Volume percentage distribution of ejected volume to the total ventricle volume for SRV patients and LV patients.

The graph in Figure 2 right shows the volume percentage distribution of ejected volume to the total ventricle volume for SRV patients and LV patients. A mean value curve is plotted for the SRV and LV patient group.

## Discussion

In this study, the flows in the pathological SRV and healthy LV were analyzed using 3D data acquired with echocardiography. For this purpose, data from five Fontan patients and two healthy patients were implemented in a moving mesh simulation model. The results of the flow simulation identified significant difference in the ventricles.

### Intraventricular flow

The investigation of blood flow within the ventricle is based on the moving mesh model, which includes the movement of the ventricular outer wall as a boundary condition. The model was used to perform patient-specific analyses of cardiac flows and to investigate the differences between SRV and LV for seven subjects.

The vortex formation in healthy ventricles compared to pathological SRV, showed a more distinct development in healthy patients. According to the literature, a main vortex forms in the middle of the ventricle up to the apex [12]. With the SRV, one vortex forms at the beginning of each of the E and A waves at the inlet, splits into several elongated vertebrae, and remains in the middle of the ventricle until the end of systole. Blood flows towards the apex and towards the ventricular wall only up to half of the ventricle, which was also observed in Lampropoulos et al. [13]. Vortex formation was lower in the SRV compared to the LV.

EF is one of the basic evaluation factors in echocardiography for cardiac function [14]. Regarding the SRV patients, the mean EF was 46.4%, and of the patients studied, two had an EF greater than 50%. In the clinic, a value above 50% is taken as an indication of a patient considered healthy [15]. However, the flow simulations conducted in this study depict a significantly deficient cardiac output for SRV patients.

VFT is a dimensionless factor that has been used in several studies as an indication of an optimally forming vortex during diastole. Healthy patients have a VFT around 4 [12]. A significantly lower value indicates a weak diastolic filling phase. The examined SRV had VFT values of 1.16 ± 0.53. This suggests a weak diastolic vortex. This vortex stores kinetic energy, which initiates the ejection of blood during late diastasis. The formation of a weak diastolic vortex can also be observed in the examined TKE. SRV patients have significantly lower TKE than the compared patients with healthy ventricles. Optimal filling and formation of TKE during diastole is necessary for sufficient energy for ventricular ejection during systole [13].

The washout simulations highlight the weakly forming vortex during diastole and the related insufficient filling of the ventricle. It was observed that residual blood remains longer in the apex of the SRV compared to the LV. After three cycles, SRV patients have an average residual volume of 24% in the ventricle. Healthy patients with LV have a residual volume between 5 and 7% after three cycles [12]. The washout simulations in this study are able to provide a clear representation regarding the performance of a ventricle.

During the study, a counterintuitive relation between TKE and VFT was observed in two SRV patients (#1, #3). These patients showed an increased TKE (2.178 J/kg; 1.041 J/kg) compared to the average value (0.881 J/kg), which indicates good ventricular performance. However, the VFT (0.62; 0.62) for these patients was below the mean value (1.16), indicating a decreased diastole and therefore a weakened performance. In parallel, an increased systolic TKE to total cycle recorded TKE ratio (75.2%, 73.4%) to the mean (54.3%) was observed. This may suggest that in the SRV patients the weak diastole is compensated with an increased systole. This relationship should be further investigated in future work.

### Moving mesh methodology and handling of the geometrical data

The advantage of the acquisition technique with the iE33 Philips ultrasound scanner is the possibility of patient-friendly and time-discrete imaging of a cardiac cycle. 3D echocardiography is a non-invasive and reliable technique that can be used at the bedside. Disadvantages of this imaging technique (related to its application in CFD simulations with a moving mesh methodology) are on the one hand the quality of the outer mesh for the geometric form. During ventricular contraction, the triangular mesh, which is constant for all time steps, causes strong deformations of the individual cells. This can lead to distorted cell nodes or illogical shapes. A manual editing of the geometry outer surface allows to obtain feasible mesh models for numerical analysis, but is time-consuming. On the other hand the impossibility of the imaging of the mitral and aortic valve shows up a second disadvantage. Here additional data with e.g. transesophageal echocardiography might be an imaging modality for good resolution of the valve structured.

### Limitations and outlook

Due to the limited field of view of the ultrasound images and the insufficient possibility of evaluating clinical 3D-data using the software ImageArena, one of the most important limitations was the lack of information about the valves and the atrium. The dynamics of the valve and the geometry of the atrium affect the specific properties of the inflow jet [9]. To reduce further inaccuracies of spatial description and low time resolution, parameters were integrated primarily over the entire ventricular volume and cardiac cycles.

Another challenge in the evaluation of univenticular hearts is the inhomogeneous Fontan patient population. The complexity of the pathology present and the surgical history of these patients is often highly individualized. Because of this limitation, we have focused only on the anatomy of singular right ventricles in this study for the time being.

The study was limited by the small amount of the analyzed patients. The limited number of the patient group resulted from the poor quality of the acquired 3D models and the time-consuming processing of them. Nevertheless statistical significant results could be observed for washout, TKE and VFT.

Further work should focus on evaluating and comparing a greater number of patients with univentricular hearts and healthy hearts. That way, the correlations observed in this work could be further quantified. The complex patient history of Fontan patients and the related anatomical conditions must always be kept in mind.

Furthermore, it is important to integrate a directed flow into the simulation model. Preferably under integration of heart valves. Subsequently, the simulation models require a qualified validation. This could be done in the form of a suitable PIV study.

### Conclusion

Looking at the EF of SRV, the cardiac work could be assumed to be moderate, but the comparison with the numerical analysis shows otherwise. Especially the washout is a clear indicator.

This study has shown significant differences in intraventricular flow between healthy patients and Fontan patients. Weak vortex formation during diastole is characteristic for the examined SRV patients. This correlation is also evident in the significantly lower TKE and in the weaker washout.

It should be considered that the studied patients have an average EF of 46%, indicating only moderately impaired cardiac conduction, but according to our washout simulations up to 20% less blood is transported through the ventricle compared to the healthy ventricle. This relationship and its relevance to the clinic should be further investigated.

## Conflict of Interest Statement

All authors have nothing to disclose.

## Author contributions

AG designed the study, performed and evaluated the simulations. JK and NW supported the method development and the conductance and analysis of the simulations. CW, KL and UH provided the clinical data, the segmented ventricular shapes and provided support during the analysis of the results. SGH and UST supported the study design and evaluation. MIN provided guidance on the analysis and interpretation of the results, supervised the study design and execution. The interpretations of the study were made jointly by all co-authors. AG wrote the manuscript based on the input of the co-authors.

## Acknowledgements

This study was funded by Fördergemeinschaft Deutsche Kinderherzen e.V. – PN: W-BN-011/2017 KH BN. MN acknowledges financial support through the ‘‘Return Fellowship” of the German Research Foundation (DFG).

## Notes

### Competing Interest Statement

The authors have declared no competing interest.

## References

[1] Krishnan U. Univentricular heart: management options. Indian journal of pediatrics 2005; 72: 519–524.

[2] Hoffman JIE, Kaplan S. The incidence of congenital heart disease. Journal of the American College of Cardiology 2002; 39: 1890–1900.

[3] Borth-Bruhns T, Eichler A. Pädiatrische Kardiologie. Berlin, Heidelberg, s.l.: Springer Berlin Heidelberg 2004.

[4] Davies RR, Chen JM, Mosca RS. The Fontan Procedure: Evolution in Technique; Attendant Imperfections and Transplantation for “Failure”. Seminars in Thoracic and Cardiovascular Surgery: Pediatric Cardiac Surgery Annual 2011; 14: 55–66.

[5] Pedrizzetti G, La Canna G, Alfieri O, Tonti G. The vortex--an early predictor of cardiovascular outcome? Nature reviews. Cardiology 2014; 11: 545–553.

[6] Mao W, Caballero A, McKay R, Primiano C, Sun W. Fully-coupled fluid-structure interaction simulation of the aortic and mitral valves in a realistic 3D left ventricle model. PloS one 2017; 12: e0184729.

[7] Chen L-J, Tong Z-R, Wang Q, Zhang Y-Q, Liu J-L. Feasibility of Computational Fluid Dynamics for Evaluating the Intraventricular Hemodynamics in Single Right Ventricle Based on Echocardiographic Images. BioMed research international 2018; 2018: 1042038.

[8] Ballyk PD, Steinman DA, Ethier CR. Simulation of non-Newtonian blood flow in an end-to-side anastomosis. Biorheology 1994; 31: 565–586.

[9] Seo JH, Vedula V, Abraham T, et al. Effect of the mitral valve on diastolic flow patterns. Physics of Fluids 2014; 26: 121901.

[10] Wilcox DC. Turbulence modeling for CFD. 2nd ed. La Cañada, Calif.: DCW Industries 1998.

[11] Sonntag SJ, Kaufmann TAS, Büsen MR, et al. Numerical washout study of a pulsatile total artificial heart. The International journal of artificial organs 2014; 37: 241–252.

[12] Mangual JO, Kraigher-Krainer E, Luca A de, et al. Comparative numerical study on left ventricular fluid dynamics after dilated cardiomyopathy. Journal of biomechanics 2013; 46: 1611–1617.

[13] Lampropoulos K, Budts W, van de Bruaene A, Troost E, van Melle JP. Visualization of the intracavitary blood flow in systemic ventricles of Fontan patients by contrast echocardiography using particle image velocimetry. Cardiovascular ultrasound 2012; 10: 5.

[14] Mor-Avi V, Lang RM, Badano LP, et al. Current and evolving echocardiographic techniques for the quantitative evaluation of cardiac mechanics: ASE/EAE consensus statement on methodology and indications endorsed by the Japanese Society of Echocardiography. European journal of echocardiography: the journal of the Working Group on Echocardiography of the European Society of Cardiology 2011; 12: 167–205.

[15] Kumar V, Abbas AK, Fausto N, Aster JC. Robbins and Cotran Pathologic Basis of Disease, Professional Edition E-Book. 8th ed.: Saunders 2009.

